# Adjustments to the reference dataset design improves cell type label transfer

**DOI:** 10.1101/2023.01.25.525533

**Authors:** Carla Mölbert, Laleh Haghverdi

## Abstract

The transfer of cell type labels from prior annotated (reference) to newly collected data is an important task in single-cell data analysis. As the number of publicly available annotated datasets which can be used as a reference, as well as the number of computational methods for cell type label transfer are constantly growing, rationals to understand and decide which reference design and which method to use for a particular query dataset is needed. Here, we benchmark a set of five popular cell type annotation methods, study the performance on different cell types and highlight the importance of the design of the reference data (number of cell samples for each cell type, inclusion of multiple datasets in one reference, gene set selection, etc.) for more reliable predictions.

## 1 Introduction

Identification of cell types is an essential part of the analysis of single-cell RNA-seq data, and provides thorough summarizing of the data in light of already existing biological context for the known cell types. Yet, often this is not a straight-forward part of processing and careful cell type annotation is a time consuming process. Recently, more attention has been devoted to development of methods for cell type annotation based on label transfer from previously annotated dataset. Several label transfer approaches have been proposed, based on different models such as correlation between the cell states (e.g. Seurat [Stuart et al., 2019], SingleR [Aran et al., 2019], CellID [Cortal et al., 2021]), random forest (e.g. SingleCellNet [Tan and Cahan, 2019])), or deep learning (e.g. ItClust [Hu et al., 2020]). Existing methods often perform well in predicting cell types of distinct clusters, while cell types without a clear boundary between them (e.g. as in continuous developmental trajectories) are more difficult to identify. Reliable annotation of rare cell types is also an important challenge and implies that the prediction quality per cell type needs to be assessed rather than reporting only overall statistics which miss any indication on where (and why) the prediction errors take place.

Here, we use reference and query scRNA-seq datasets from the Peripheral Blood Mononuclear Cells (PBMC) samples [Ding et al., 2020] to benchmark five popular label transfer methods and show that the design of the reference dataset should be adapted to the learning approach used by the method. We use reference and query datasets which are complete with respect to each other, meaning that all the cell types in the query are present in the reference and vice versa. First, we examine the effects of reference data sampling (i.e., number of cells per cell type) on each method. Using a balance training set is standard in the machine learning field [Kotsiantis et al., 2005] but has mostly been neglected for cell type annotation. We then, introduce a weighed bootstrapping-based approach to make use of as much of the reference data as possible, while still keep the benefits of having a balanced reference dataset. We further show the effect of using reference data from various sources on the different methods, that careful selection of the gene set is crucial for the quality of label predictions for high-dimensional and noisy scRNA-seq data and that high confidence scores provided by the different methods do not directly correlate with correct predictions.

## 2 RESULTS

### 2.1 Less abundant cell types are more difficult to predict

We evaluate the methods on a PBMC dataset, with manually curated cell type labels (Figure 1A). We start by using the entire reference data without adjustment. Comparing each method’s predictions to the ground truth, we get similar prediction accuracy, except for CellID performing significantly worse (Figure 1B). Comparing the ground truth with the annotation from each method, we see that, even though the accuracy is similar for most methods, the areas of difficulty vary (Figure 1C and D). In general the mispredictions are located in areas of the Uniform Manifold Approximation and Projection (UMAP) [McInnes et al., 2018, McInnes et al., 2020] where cell types overlap. Taking for example the area where the two rare cell types of Dendritic cells and Megakaryocytes overlap in the ground truth, we see different predictions with each method. Seurat extends the B cell cluster that lies below, while SingleCellNet predicts a mixture of different cell types in that area. SingleR on the other hand predicts the Dendritic cells in a much wider radius then the ground truth. ItClust does not predict any cells as either Dendritic cells or Megacaryocytes, thereby completely mispredcting these populations. This shows, that even though these four methods have similar overall accuracy the prediction for specific populations can vary greatly. This variation is highlighted by the cell type specific accuracy and precision values for each method. All methods perform worse for rare cell types as reflected in decrease of accuracy or precision (figure 1E and F). ItClust even misses four of the less abundant cell types completely.

**Figure 1:**
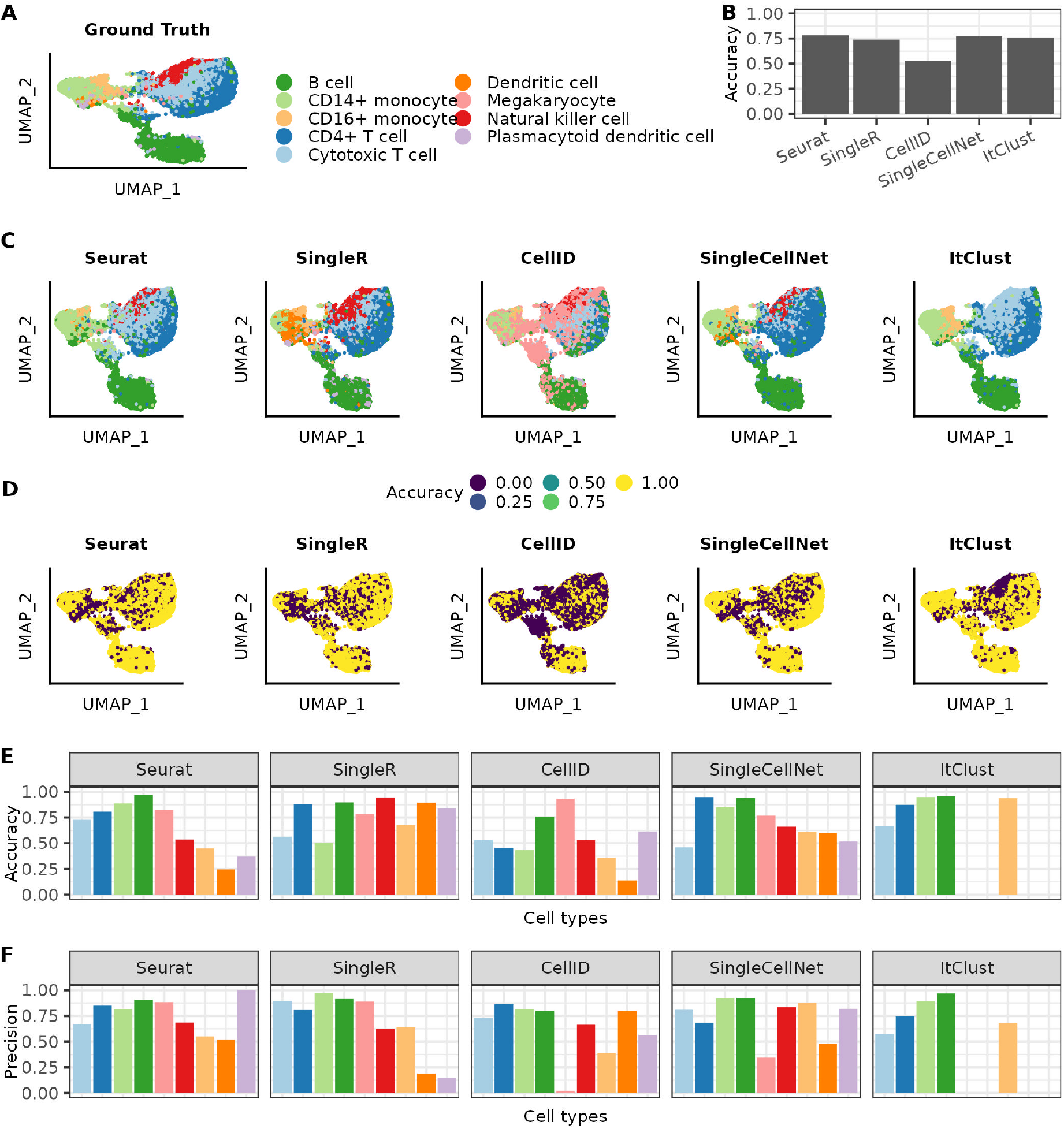
Cell type label transfer on the PBMC dataset using the full reference dataset. (A) UMAP colored based on the cell types assigned in the ground truth. (B) Over all accuracy achieved for each of the methods. (C) UMAPs colored by the cell type annotation made with the different annotation methods. (D) UMAPs colored by if a cell was correct or incorrect predicted. (E) Accuracy and (F) Precision achieved for the different cell types with each of the different methods. The cell types are listed in decreasing order of how often they are represented in the full reference data.

### 2.2 Less abundant cell types benefit from more balanced reference data

Since rare cell types are more difficult to predict, we asses how a more balanced reference dataset can affect the predictions. We compare the accuracy of the predictions for each cell type (three example cell types in Figure 2A and the remaining cell types in S1) as a function of the maximum number of cells per cell type in the reference dataset. There are no visible improvements when increasing the number of cells per cell type to more than 1000 cells, but especially for less abundant cell types it can lead to a drop in accuracy. This decrease is especially drastic for ItClust where smaller cell types start to be missed completely. On the other hand including to few cells per cell type has negative effects on the predictions, especially for abundant cell types. When comparing the precision of the predictions as a function of the maximum number of cells per cell type (Three example cell types in Figure 2B and the remaining cell types in S2) we see that it is less dependent on cell type balance in the reference data. Overall the quality of the predictions for each cell type depends on how well they are represented in the reference data. Each cell type tends to be predicted best, when maximum number of cells per cell type is closest to the maximum number of cells for that cell type. At this point the cell type is best represented without being overshadowed by other cell types.

**Figure 2:**
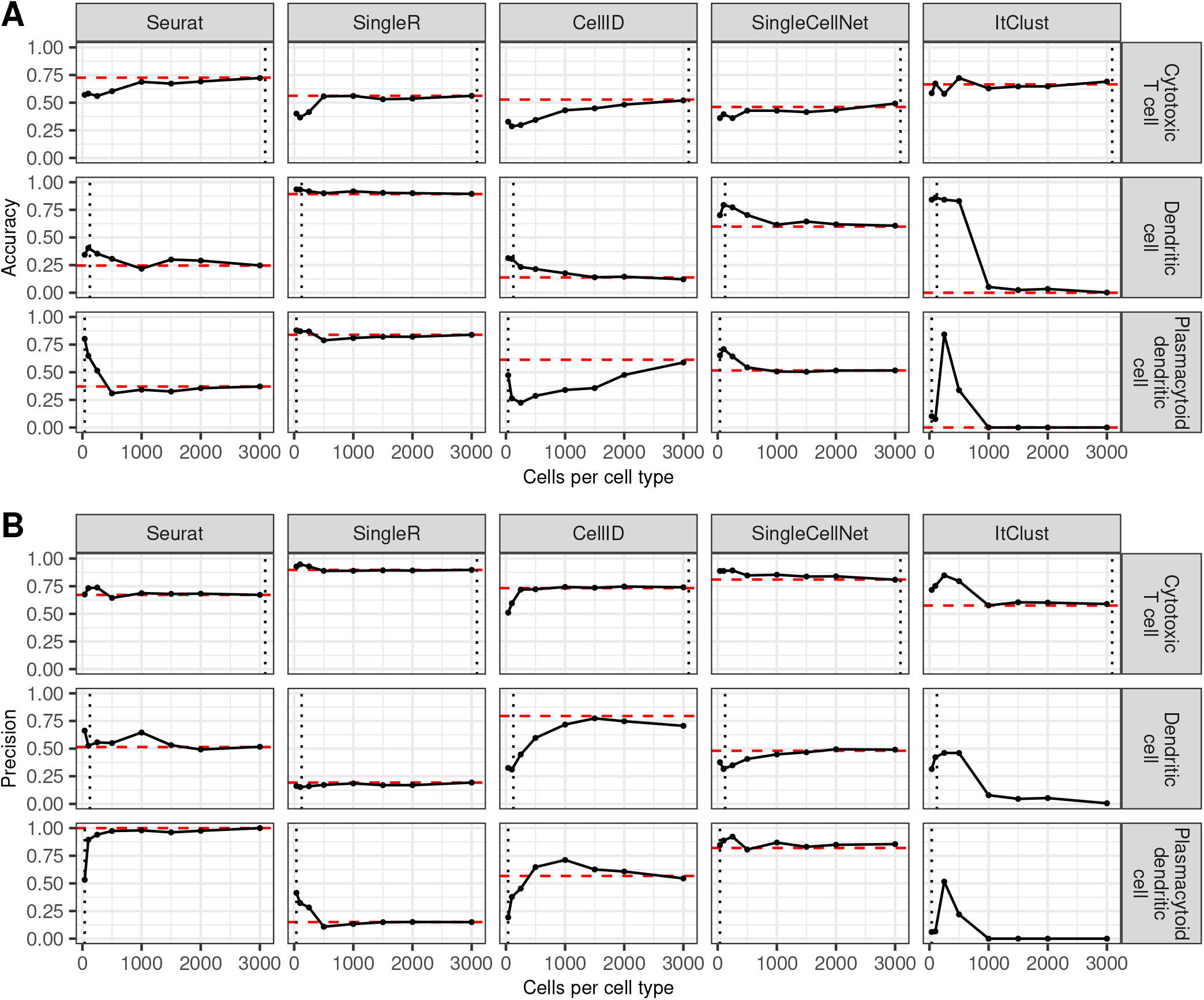
Effect of the number of cells per cell type on the predictions. Distribution of the accuracy (A) and precision (B) for three example cell types in each method, when the number of cells per cell type is increased. The red line shows the accuracy on the full data and the grey line shows the number of cells in this cell type in the full reference data. For the remaining cell types see supplementary figures S1 and S2

### 2.3 Weighted bootstrapping increases the accuracy in the prediction of less abundant cell types

Here, we introduce a weighted bootstrapping-based approach, that allows us to account for the abundances of each cell type in the reference data (see methods 4.4). This allows us to use as much of the reference data as possible, without limiting ourselves to a balanced subset. Figure 3A shows how well each cell is predicted across 20 bootstrapping sets. We observe that the lowest accuracy of each method mirrors the results shown in figure 1D. Figure 3B and C summarizes the performance for each cell type in accuracy and precision. With bootstrapping the accuracy increases for all smaller cell types but their precision tends to decrease. This is especially visible for Seurat and SingleCellNet. SingleR is not much affected by the bootstrapping, neither positive nor negative. Overall bootstrapping does not improve the predictions of CellID. ItClust performs better for all cell types with the additional advantage that no cell types are missed completely.

**Figure 3:**
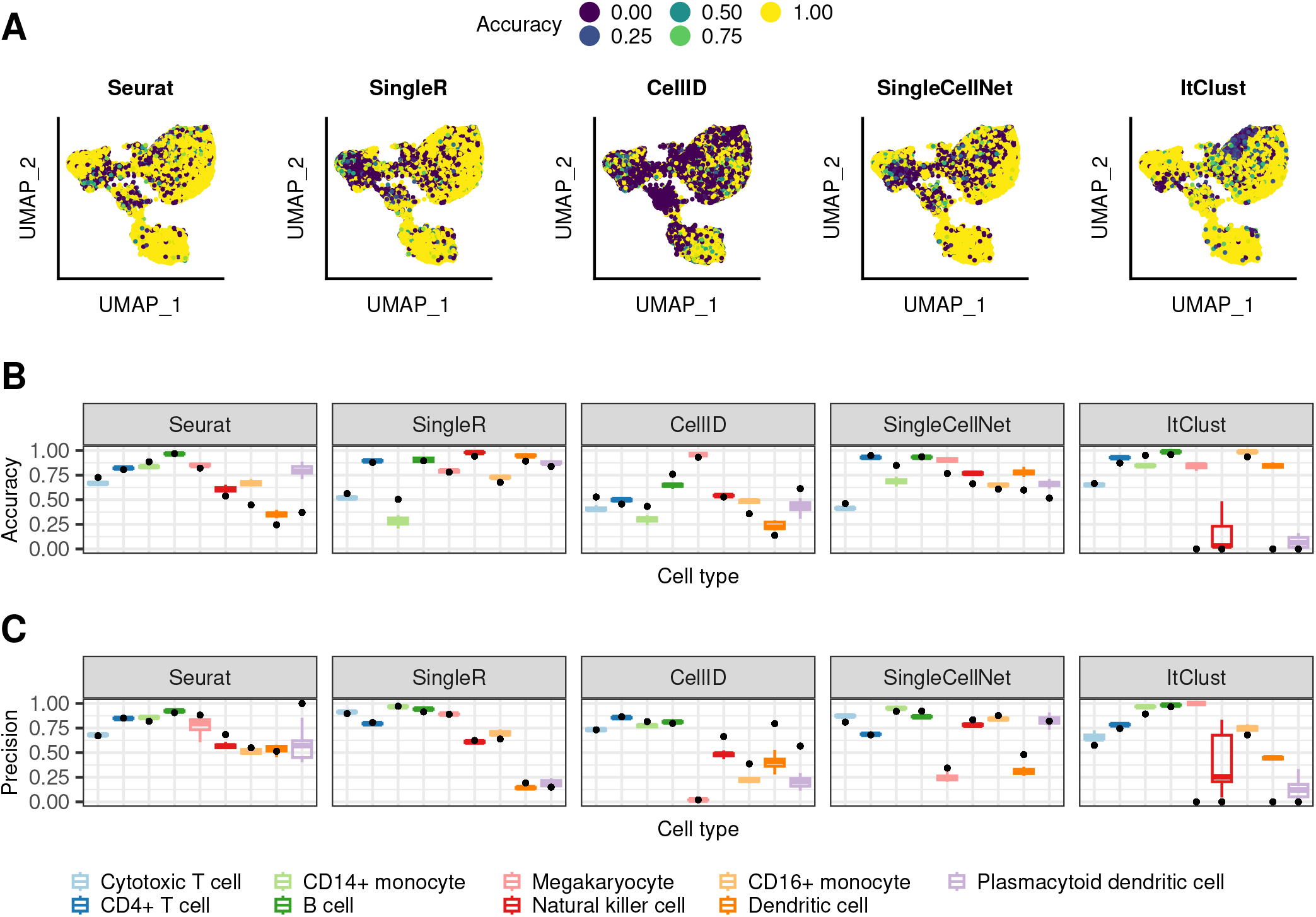
Using weighted bootstrapping for the predictions. (A) UMAP colored by the difference between the mean accuracy achieved when using the bootstrapping-based approach versus the one achieved on the individual reference subsets. (B) Distribution of the accuracy (B) and precision(C) for the individual reference subsets shown as boxplots (see Methods). The black points represents the result achieved on the full reference set.

### 2.4 Including data from different sources affects the methods differently

With the increasing number of annotated datasets, it becomes possible to combine multiple existing datasets into the reference. To simulate this we extended our reference with cells from other sequencing technologies and repeated the annotation using the bootstrapping approach. The UMAPs colored by the difference in accuracy for each cell when using a mosaic-reference set compared to a mono-source reference set, shows that most cells have a similar performance for both sets and that changes mostly occur in difficult areas (figure 4A). Taking a closer look at the cell type specific accuracy and precision values, the use of mono-source data has no significant effect on the abundant cell types in any of the methods (figure 4B and C). For Seurat, SingleR and ItClust the performance on some of the rare cell types is improved while it gets worse on others, making it difficult to show a clear benefit or deficit from using mosaic-data. CellID and SingleCellNet do not decrease in performance for any of the cell types but some smaller cell types are predicted better, showing that here mosaic data is beneficial.

**Figure 4:**
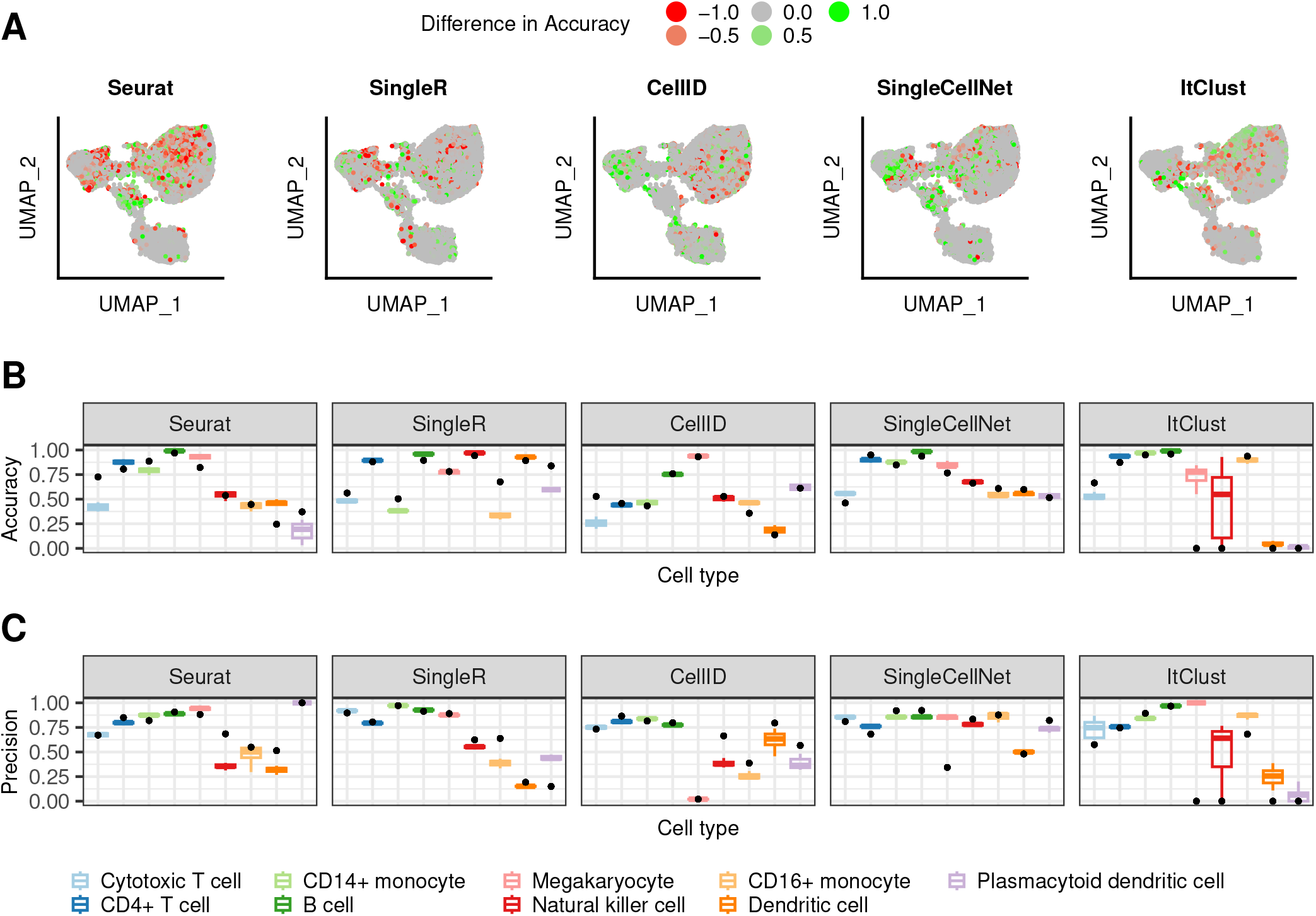
Using mosaic reference data versus the mono-source reference. (A) UMAP colored by the mean accuracy achieved on the mosaic references subtracted by the mean accuracy achieved on the mono-source references when using the weighted bootstrapping approach (B) Distribution of the accuracy (B) and precision (C) for the mosaic (colored boxplots, see Methods) for the weighted bootstrapping-based approach. The initial performance on the full refnerence is shown as a comparison (black dot).

### 2.5 Selection of the gene set affects the methods differently

In the previous sections we used a set of 200 highly variable genes (HVGs) for the predictions. To test whether using a different gene set would affect the predictions by the different methods, we reran all the methods for 1000 HVGs. In figure 5 we show the accuracy and precision achieved when using either 200 or 1000 HVGs. While the gene set affects the performance, the results differ significantly between the different methods. Seurat appears to benefit from more HVGs, when it comes to the prediction of less abundant cell types. For SingleR the less abundant cell types benefit from more HVGs, when it comes to the precision of the predictions. But abundant cell types benefit from keeping a low number of HVGs. Increasing the number of HVGs would lead to an accuracy below 0.5 for two of the more abundant cell types. CellID shows the most severe difference in accuracy and prediction, when it comes the HVG selection. Using 1000 HVGs leads to almost all cells being incorrectly predicted. SingleCellNet performs slightly better in regards to the accuracy when including 1000 HVGs, but the precision is significantly better when limiting the genes to the top 200 HVGs. ItClust includes internal filtering of the gene set. Therefore was excluded for the analysis in this section.

**Figure 5:**
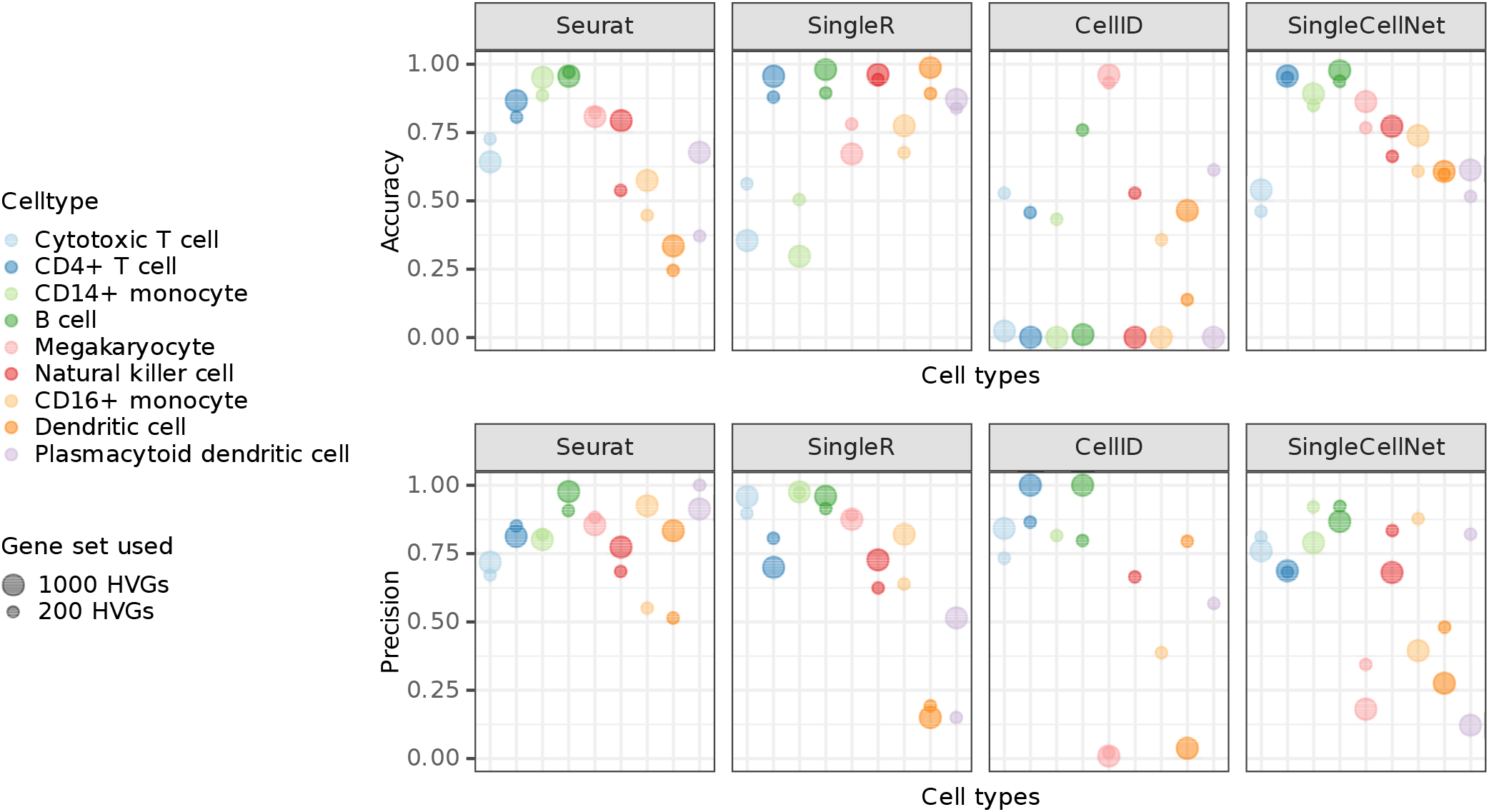
Comparing predictions based on two different gene sets. (A) Cell type specific accuracy (A) and precision (B) achieved with the different gene sets for each of the methods.

### 2.6 High confidence scores do not always align with correct predictions

Four of the five methods of interest supply a confidence score for the predicted cell type. In figure 6A and supplementary figure S3 we show the confidence scores each of the methods provided separated in true and false predictions. While Seurat shows for many cell types a clear difference in distribution of confidence scores, for true and false predictions, there are cell types where this difference is rather small, such as the Cytotoxic T cells and the natural killer cells. For less abundant cell types such as the Dendritic cells and the Plasmacytoid dendritic cells, the true predictions have generally a higher confidence than the false predictions, for smaller numbers of cells per cell type. But with an increase in the number of cells, Seurat starts to become even more confident in the false predictions, making the confidence scores less helpful. This implies, the method does not model the rare cell types correctly and only becomes more confident in a false model for them when provided larger number of cells which make the data more imbalanced. The confidence scores supplied by SingleR show significantly higher confidence for correct predictions in the rare cell types (non-overlappling error bars for correct and false predictions indicate a significant difference between them), but no clear difference between the two distributions for some other cell types e.g., Cytotoxic T cells. This implies a good modeling of rare cell types, but again not a good model for cell types with mixing boundaries. Similarly, for SingleCellNet the confidence in the correct predictions is generally (but not for all cell types) higher than the one in false predictions, which is good. For ItClust, the confidence scores appear as rather random and uninformative. The (partial) discordance we observe between confidence scores and correctness of predictions across the methods is not surprising, as the confidence score reflects the variation in prediction for a query cell given a model; when the model is wrong (or not good enough), it can make wrong predictions quite confidently. CellID does not provide confidence scores, so it was excluded for the analysis in this section.

**Figure 6:**
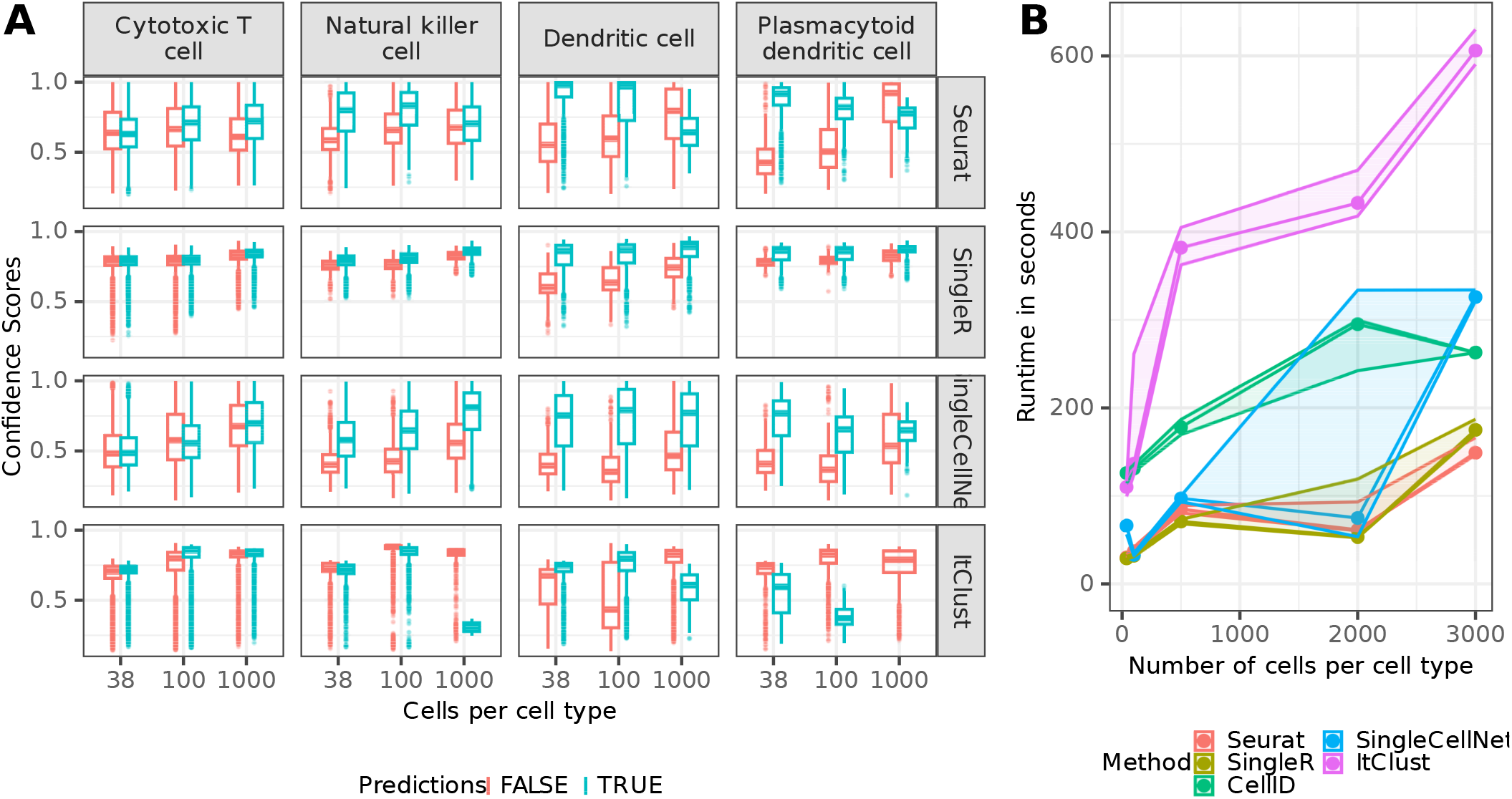
Confidence scores and runtime. (A) Boxplots (see Methods) of confidence-scores for the predicted cell types depending of true and false predictions of each of the methods providing confidence scores for three example cell types. For the other cell types see supplementary figure S3. (B) runtime in seconds on a CentOS7 cluster with a memory limit of 16G on one core with 8CPU available.

### 2.7 Reporting the runtimes

Additionally, to the accuracy of the different methods we looked at their runtime (Figure 6B), which revealed that runtime of all methods increases linearly or slower with the number of cells in the reference. For all trials the query data was fixed. Seurat and SingleR are the two most efficient methods and ItClust is taking longer than all the other methods even with small reference datasets.

## 3 DISCUSSION

In this work we benchmarked five popular label transfer methods on PBMC datasets and showed how the selection and treatment of the reference data affects the quality of the prediction. All methods tend to predict highly represented cell types better than rarer cell types. Reducing the reference data to balance the cell types improves each methods ability to predict rarer cell types. However, this comes at the cost of lower accuracy of abundant cell types. To account for this, we introduce a weighted bootstrapping method that includes the predictions of multiple different sized subsets. This method generally improves the prediction accuracy for rare cell types, but this can come at the cost of precision of these cell types. Overall methods that include an explicit modelling step (different sorts of data compression, deep learning, etc) benefit from more balanced reference sets, in contrast to methods that rely only on cell states’ correlations such as SingleR.

The number of available annotated datasets is continuously growing and combining multiple datasets could potentially increase the statistical power of data integration due to increased data size. However, multiple factors such as technical differences and batch effects between the datasets can introduce new causes for mispredictions. To assess this, we combined reference datasets from multiple sources (without batch correction) and evaluated how this affected the predictions. Here again the answer depends on the annotation method. For CellID and SingleCellNet the mosaic reference lead to a clear improve in predictions, while the benefits are less significant for the other methods. In general mosaic data does not appear to weaken the overall performance for any method. Thus, especially in cases where combining multiple reference datasets would allow identification of additional cell types, we would suggest to do so. Also in rare cell types, such as the Plasmacytoid dendritic cells for which the minimum number of cells is increased from 38 in the mono-source to 102 in the mosaic data, the addition of data appears to be beneficial for all correlation based methods (Seurat, SingleR, CellID).

When determining distances between different cell states in high-dimensional space, we often face the challenge of the curse of dimensionality [Imoto et al., 2022]. The higher the number of dimensions, the more severe the issue, as noise disproportionately adds up to undermine the true (biological) signal when considering multiple dimensions. As such, one could expect that reducing the number of dimensions helps with better defining cell similarities (and distances), thus improving the predictions. In a previous study [Triana et al., 2021], using a curated set of 462 genes with known biological relevance facilitated reliable cell type label predictions for a bone marrow datasets. However, such prior knowledge may not be at hand for the data analyst. Without using prior knowledge, we compared performances between two gene sets of different sizes (200 and 1000 top HVGs) and showed that the number of included HVGs alone is not determinant of the predictions quality. In fact using different gene sets affected different cell types and methods differently. For Seurat and SingleR that already perform well when using as little as the top 200 HVGs, predictions for most cell types were improved by including more HVGs. These methods in fact, have their own inner procedure for reducing the number of dimensions (see Methods). For the random forest-based method SingleCellNet higher accuracies when using the larger gene set, come on the cost of lower precision.

We demonstrated that confidence scores provided by different methods, cannot be taken as an absolute proxy for correctness of predictions, but rather they show the robustness of predictions assuming the model used by the method is correct. Therefore, a correlation between the confidence scores and correct predictions (when ground truth labels are available) would indicate the reliability of a model. Overall, SingleR showed one of the most robust and reliable performances in our benchmarking experiments using PBMC reference and query datasets that were complete with respect to each other. Seurat and SingleCellNet’s performances were also generally good and competing with SingleR, but we assessed CellID and ItClust as not being very robust and reliable. However, one could expect the performances to be different for other data scenarios, which thus need to be further tested, e.g. scenarios in which one or a few cell types are exclusively present in either the query or the reference set, or when much bigger reference data is available. In our study, none of the available methods were able to make highly accurate predictions for all cell types even with careful design of the reference. This could be due to inadequate methods’ performance, but also to incorrectly labeled cells in the reference and query that we assume as ground truth. The annotations assumed as ground truth in this study have been attained in the original publication [Ding et al., 2020] using standard single-cells data clustering and annotation algorithms, which may include errors. As figure 1 indicates, annotation errors as well tend to happen more along the boundaries between closely related cell types (e.g., in continual developmental trajectories). The definition of these boundaries diverges in different annotations, which makes some degree of error in such regions inevitable.

To conclude, our study highlights the need for further improvement of cell type annotation and label transfer methods as well as better data quality acquisition. With restriction to the existing methods, we showed that different methods’ predictions can be improved by a careful design of the reference. In particular, this design needs to be adapted to the method used.

## 4 MATERIALS AND METHODS

### 4.1 Data availability

We performed our analysis on publicly available human PBMC cells [Ding et al., 2020] data (table 1, Gene Expression Omnibus accession number: GSE132044). The data has been annotated in the original publication, which we use as the ground truth cell labels going forward. The code for reproducing the results and figures in this study is available on GitHub (https://github.com/HaghverdiLab/cellAnnotation_Benchmark).

**Table 1:** Description of all datasets included in this study

| Dataset | No. cells | No. genes | No. classes | Description | Protocol |
| --- | --- | --- | --- | --- | --- |
| PBMC Query | 11183 | 33658 | 9 | PBMC | Smart-Seq2, CEL-Seq2, 10X, Drop-Seq, Seq-Well, inDrops |
| PBMC 10X | 9666 | 33658 | 9 | PBMC | 10X |
| PBMC CEL-Seq | 253 | 33658 | 7 | PBMC | CEL-Seq2 |
| PBMC Drop-Seq | 3222 | 33658 | 9 | PBMC | Drop-Seq |
| PBMC inDrops | 3222 | 33658 | 7 | PBMC | inDrops |
| PBMC Seq-Well | 3176 | 33658 | 7 | PBMC | Seq-Well |
| PBMC SMART | 253 | 33658 | 6 | PBMC | Smart-Seq2 |

### 4.2 Preprocessing

The PBMC sample in [Ding et al., 2020] contain two experiments run on different days. Both experiments contain samples from all scRNA-seq platforms. In sections 2.1 to 2.3 and 2.5 to 2.7 we use exclusively the cells in “experiment one” which were gathered on the 10X platform as our reference data, which we refer to as the mono-source reference. In section 2.4 we combine the 10X data with all the other scRNA-seq platforms (i.e., Smart-Seq, CEL-Seq2, 10X, Drop-Seq, Seq-Well, inDrops) from the same experiment into one reference, which we refer to as the mosaic reference. We use experiment two as query data. The query and the reference data contain the same nine cell types (Cytotoxic T cellm CD4+ T cell, CD14+ monocyte, B cellm Megakaryocyte, Natural killer cell, CD16+ monocyte, Dendritic cell, Plasmacytoid dendritic cell).

In order to make the different label transfer methods comparable, we use the same data preprocessing workflow for all methods as far as possible. The preprocessing of the gene expression counts data starts with selecting the top 200 HVGs (or 1000 HVGs) using the Pearson residuals method [Dmitry Kobak, 2020]. We then do a log transformation of the highly variable genes (HVGs) read counts followed by cell-wise L2 normalization. To allow the log transform on the expression matrix, we add a small value (0.001) to the expression values in order to avoid zero values. However, these preprocessing steps could not be used for ItClust, for which the preprocessing steps are hard-coded in the package and cannot be changed. Minor adjustments to the preprocessing workflow to meet each method’s requirements are described in its corresponding Methods section.

### 4.3 Reference data permutation and confidence intervals

To analyze the importance of reference data selection, we assess the different methods, using subsets of the initial reference data of varying size. We vary the maximum numbers of cells per cell type (38, 100, 250, 500, 1000, 1500, 2000, 3000). We note that if a cell type has less then the maximum number of cell types the reference data will be unbalanced. For each of the reference set sizes, 20 random permutations were sampled from the underlying reference data.

### 4.4 Weighted bootstrap-annotation of cell types

We introduce a weighted bootstrap-annotation approach, to assign a weight to the predictions, based on how the predicted label is represented in the full reference. Each query cell *i* gets assigned a label *l_i, j_* for each reference set *r_j_*:

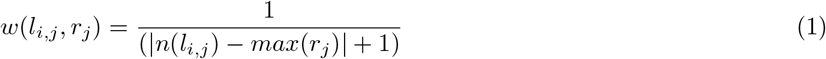

where *n*(*l_i,j_*) is the number if cells annotated with label *l_i,j_* in the full reference data and *max*(*r_j_*) is the maximum number of cells per cell type in reference set *r_j_*. For each cell type the different weights are added up and the cell type with the highest weight is predicted.

Here, we select a subset for each cell type in the reference data with the maximum number of cells per cell type being the number of cells for this cell type in the full reference.

### 4.5 Specification of the boxplots

In all boxplots in the figures of this manuscript, the box fills the interquartile range (*IQR*) between the 25th (*Q*1) and the 75th (*Q*2) percentile and the line through the box shows the median of the distribution. The lines extending from the bx show the minimum (*Q*1 – 1.5 * *IQR*) and maximum (*Q*3 + 1.5 * *IQR*) value in the data. Values outside of this range are plotted as individual points and are considered potential outliers.

### 4.6 Seurat

We follow the Seurat label transfer workflow as suggested in the [‘Mapping and annotating query datasets’](https://satijalab.org/seurat/articles/integration_mapping.html) vignette. The Seurat algorithm consists of two steps. The first is an unsupervised compression of the reference and query data into a common space that captures the most correlated features between the two datasets, by using Canonical Correlation Analysis (CCA). This step implies that the query data distribution (as well as the reference data) affects the data compression model which is used in the next step for label transfer. In the second step, the most common label among the set of mutual nearest neighbors (MNN) of a query cell [Haghverdi et al., 2018] in the reference set is transferred to it as the predicted label. We use version v4.2.0 of the Seurat R package.

### 4.7 SingleR

SingleR uses in iterative approach to transfer cell type labels from prior annotated reference data to an unannotated query dataset. The annotation process is performed individually for each query cell. First the variable genes among the cell types in the reference set are selected. Secondly, the Spearman correlation between the query cell and each cell in the reference is calculated. The correlation values are aggregated by cell type and the cell type with the lowest correlation value and all cell types with a correlation more then 0.05 smaller then the top value are removed. The steps are then repeated until only two cell types remain and the cell type with the higher correlation is predicted. We use version v2.0.0 of the SingleR R package.

### 4.8 CellID

CellID uses Multiple Correspondence Analysis (MCA) for data compression (unsupervised step) as well as to identify per-cell gene signatures. The signatures of the reference and query data are then compared using a hypergeometric test. The label of the closest cell in the reference is then transferred to the corresponding query cell. Gene set selection is part of the CellID workflow, therefore instead of initially selecting 200 HVGs, we select the top 5000 HVGs and use CellID’s specific gene set selection method to reduce the number of genes to 200. We note that this gene set varies from the one used by the other methods. We use version v1.6.0 of the CellID R package.

### 4.9 SingleCellNet

SingleCellNet is a random forest based approach, that transforms the data into a cell-by-cell binary matrix derived by pairwise comparison of selected genes (Top-Pair transformation). The workflow starts with reducing each cell type to a fixed number of cells (default: 100 cells). Here, we skip this step. This is followed by training the model. We use the default settings. The labels are transferred to the query data, based on the random forest classifier. SingleCellNet allows to set a number of random profiles to the evaluation process, which allows to identify if cells might belong to a cell type not represented in the reference data. We treat these cells as false predictions. We use version v0.4.1 of the SingleCellNet R package.

### 4.10 ItClust

ItClust is an iterative transfer learning approach for clustering and cell annotation. The neural network model is trained in two steps. It starts with supervised learning on the reference data and then additional learning step on the query data to fine-tune the parameters. We use ItClust with its default settings, which includes a gene set selection and a filtering of reference and query cells. We treat removed query cells as false predictions. Since ItClust includes its own preprocessing steps, we did not apply our preprocessing workflow. We use version v0.0.5 of the ItClust python package.

## Author Contributions

C.M. designed and performed all computational methods and benchmarking pipelines, interpreted the results and wrote the manuscript. L.H. supervised the study, contributed to the results interpretation and wrote the manuscript.

## Acknowledgments

We would like to thank Valérie Marot-Lassauzaie for providing feedback on the manuscript.

