## Supplementary Figures 1-3 for "Adjustments to the reference dataset design improves cell type label transfer"

Carla Moelbert and Laleh Haghverdi

January 25, 2023

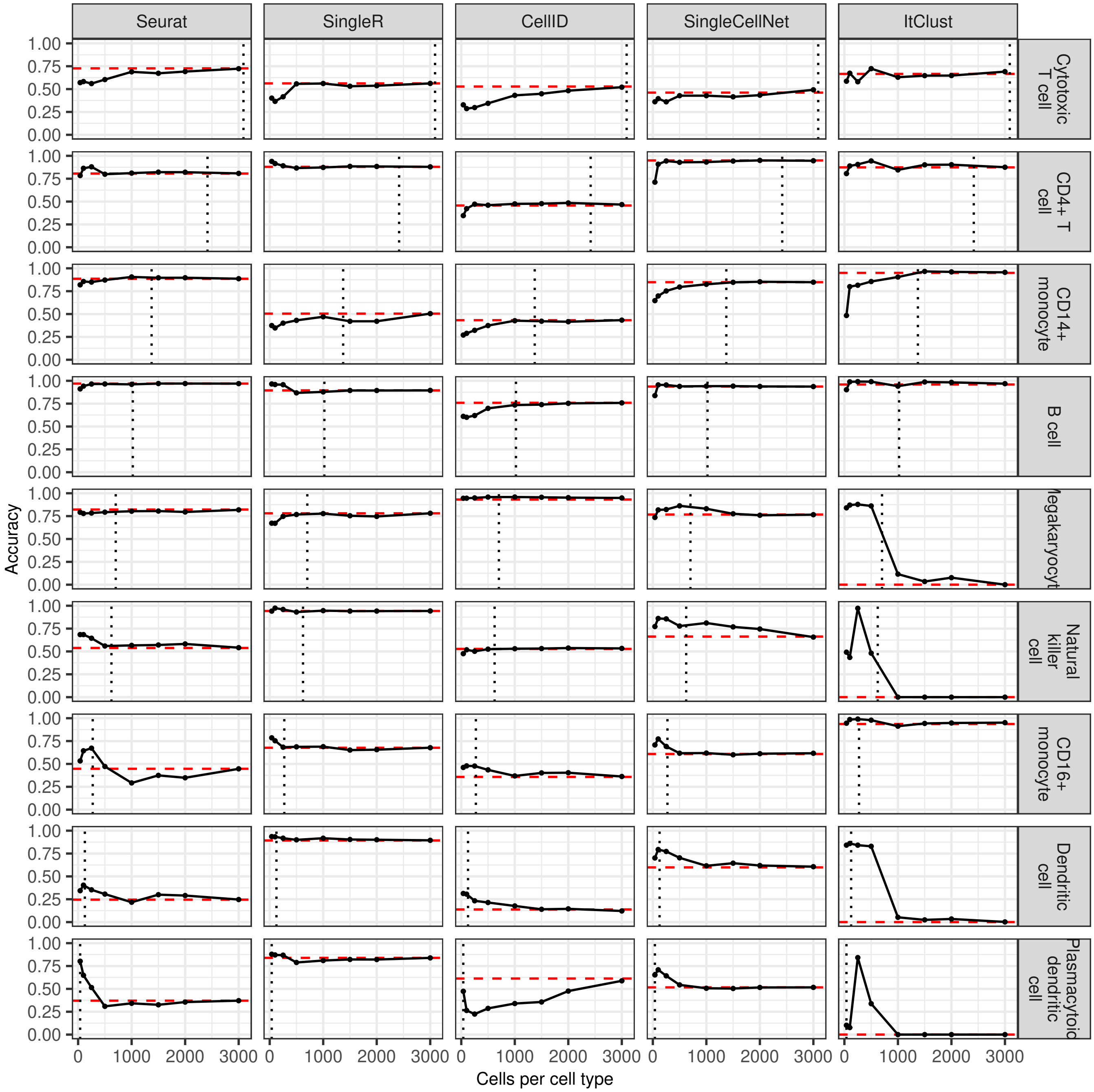


Figure 1: Distribution of the accuracy for each cell type for each method, when the number of cells per cell type is increased. The red line shows the accuracy on the full data and the grey line shows the number of cells in this cell type in the full reference data.

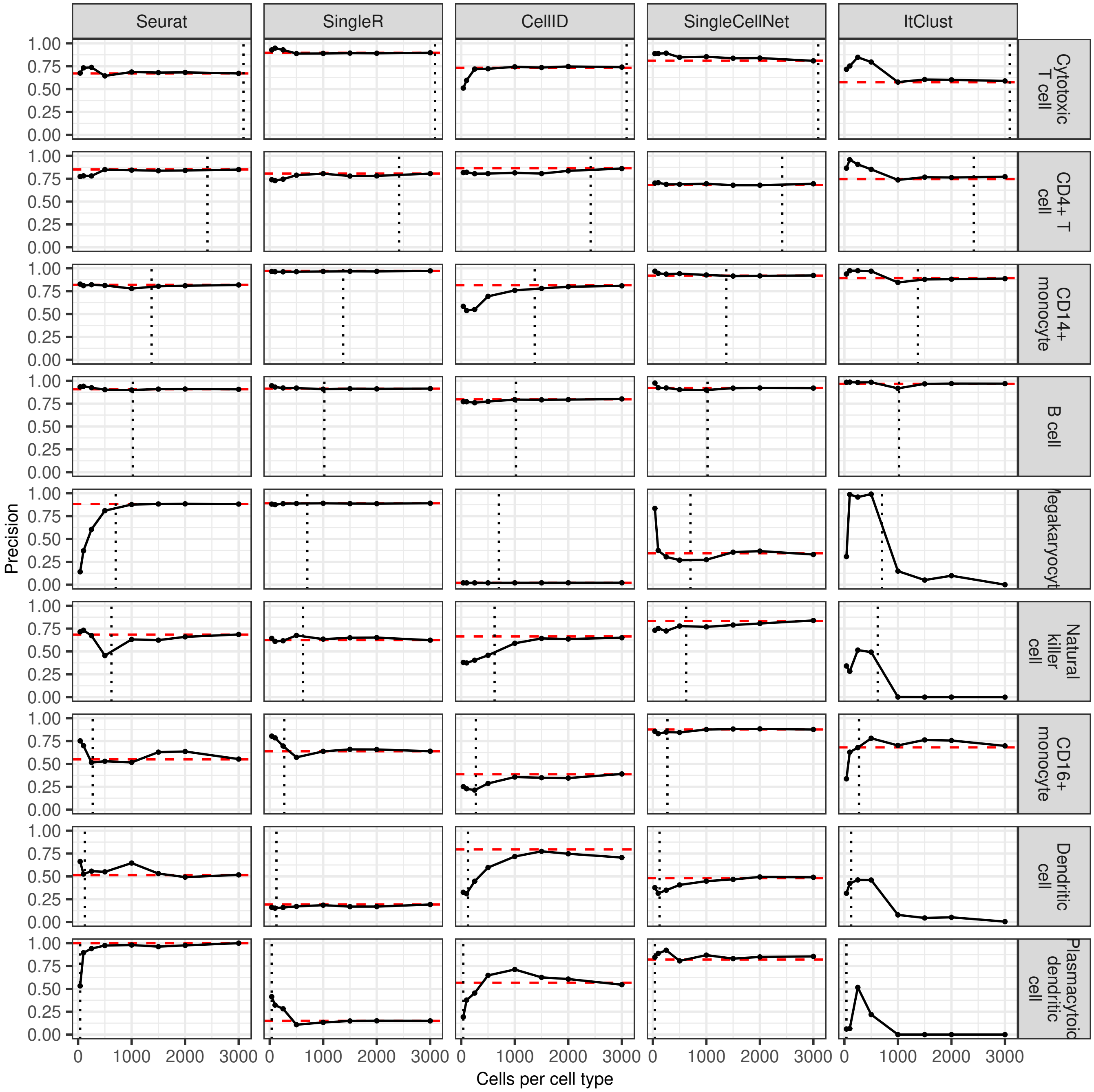


Figure 2: Distribution of the precision for each cell type for each method, when the number of cells per cell type is increased. The red line shows the accuracy on the full data and the grey line shows the number of cells in this cell type in the full reference data.

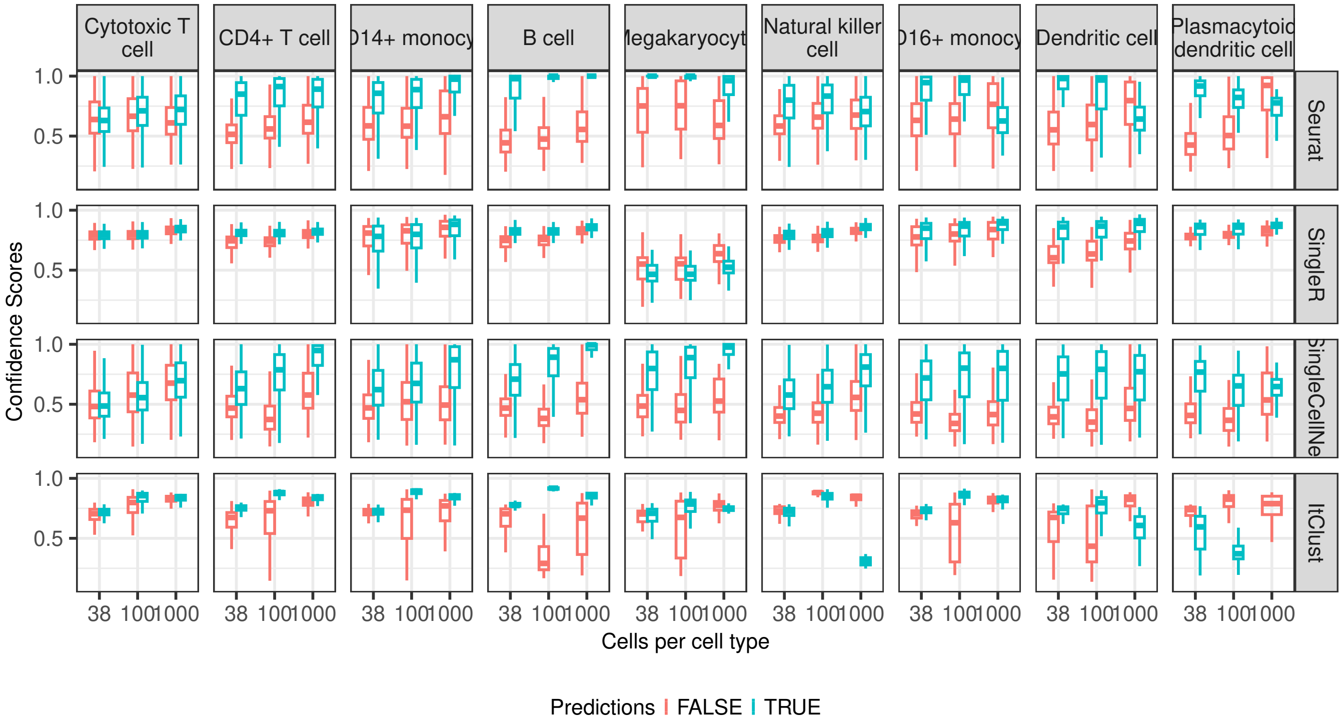


Figure 3: Distribution of confidence-scores for the predicted cell types depending of true and false predictions of each of the methods providing confidence scores.
